# Longitudinal Trajectory of Early Functional Recovery in Patients with First Episode Psychosis

**DOI:** 10.1101/525824

**Authors:** Mei-Hua Hall, Kristina M Holton, Dost Öngu□r, Debra Montrose, Matcheri S. Keshavan

**Author notes:** Address for correspondence: Mei-Hua Hall, Ph.D., Psychosis Neurobiology Laboratory, Mailstop 315, Harvard Medical School, McLean Hospital, 115 Mill Street, Belmont, MA 02478 USA.

## Abstract

**Background:** There is a large variability in the recovery trajectory and outcome of first episode of psychosis [FEP] patients. To date, individuals’ outcome trajectories at early stage of illness and potential risk factors associated with a poor outcome trajectory are largely unknown. This study aims to apply three separate predictors (positive symptoms, negative symptoms, and soft neurological signs) to identify homogeneous function outcome trajectories in patients with FEP using objective data-driven methods, and to explore the potential risk /protective factors associated with each trajectory.

**Methods:** A total of 369 first episode patients (93% antipsychotic naive) were included in the baseline assessments and followed-up at 4-8 weeks, 6 months, and 1 year. K means cluster modelling for longitudinal data (kml3d) was used to identify distinct, homogeneous clusters of functional outcome trajectories. Patients with at least 3 assessments were included in the trajectory analyses (N=129). The Scale for the Assessment of Negative Symptoms (SANS), Scale for the Assessment of Positive Symptoms (SAPS), and Neurological examination abnormalities (NEA) were used as predictors against Global Assessment of Functioning Scale (GAF).

**Results:** In each of the three predictor models, four distinct functional outcome trajectories emerged: “Poor”, “Intermediate”, “Good” and “Catch-up”. Individuals with male gender; ethnic minority status; low premorbid adjustment; low executive function/IQ, low SES, personality disorder, substance use history may be risk factors for poor recovery.

**Conclusions:** Functioning recovery in individuals with FEP is heterogeneous, although distinct recovery profiles are apparent. Data-driven trajectory analysis can facilitate better characterization of individual longitudinal patterns of functioning recovery.

## Introduction

Clinical presentations of first episode psychosis are heterogeneous (Keshavan et al., 2013). Patients with a first episode of psychosis [FEP] frequently present with a mixture of affective and psychotic symptoms, making it often difficult to make a definitive diagnosis. About a third of FEP patients receive diagnoses of affective psychosis at the end of two years and about two thirds receive a diagnosis of schizophrenia [SZ] (Keshavan and Schooler, 1992). Moreover, psychotic spectrum disorders such SZ, schizoaffective [SA], and psychotic bipolar disorder [BPD] have substantial overlap in symptoms, neurobiology, treatment response, and genetic liability (Cross-Disorder Group of the Psychiatric Genomics et al., 2013a; Cross-Disorder Group of the Psychiatric Genomics et al., 2013b; Hall and Smoller, 2009; Hall et al., 2015; Hall et al., 2013; Hall et al., 2009; Hill et al., 2013; Ivleva et al., 2012; Mathew et al., 2014; Ruderfer et al., 2014; Sklar et al., 2011; Tamminga et al., 2014). This heterogeneity is likely caused by multiple etiologies such that psychosis is likely a final common endpoint, similar to ‘congestive heart failure’ being a common endpoint of cardiac, renal and pulmonary diseases, each of which requires different optimal treatment strategies.

Psychotic disorders are accompanying by prominent social and occupational functional impairment, and represent as one of the leading causes of disability worldwide (Global Burden of Disease Study, 2015). Functional deterioration occurs at the fastest rate during the early phase of psychosis (Cannon et al., 2015; Jindal and Keshavan, 2008; Pantelis et al., 2005), which is both a “critical period” of neuronal and psychosocial plasticity and a “window of opportunity” during which treatment may confer disproportionately favorable outcomes(Caplan et al., 2013; Dazzan et al., 2015; Flyckt et al., 2006). In FEP patients, there is a large variability in the recovery trajectory and functional outcome (Allott et al., 2011; Menezes et al., 2006; Robinson et al., 2004; Strauss et al., 2010). Only 26-43% of FEP patients are estimated to achieve good global functional outcome at 1-2-year follow-up (Gonzalez-Blanch et al., 2010; Menezes et al., 2006). About 10-15% of FEP patients are highly treatment resistant. While the majority of the patients have symptomatic recovery (Verma et al., 2012) a subgroup of patients experience multiple relapses during the course of illness (Lehman et al., 2004). Also, individuals treated with antipsychotic medications may achieve good symptomatic outcomes, but have significant persistent functional disabilities, particularly in vocational and social domains (Gupta et al., 1997; Hegarty et al., 1994; Robinson et al., 2004; Tohen et al., 2000). This variability is likely due to the fact that functional recovery represents a multi-dimensional construct and many distinct domains of neurobiological alterations collectively contribute to the variance in functioning (Allott et al., 2011).

According to the literature, severity of psychopathology, particularly negative symptoms (Carlson et al., 2012; Herbener and Harrow, 2004; Ho et al., 1998; McGlashan and Fenton, 1992; Smith et al., 2002; Ventura et al., 2009), neurocognitive impairments (Best et al., 2014; Bowie et al., 2010; Green et al., 2000), and premorbid characteristics (Ayesa-Arriola et al., 2016; Bucci et al., 2016) are significant and independent predictors of functional outcome, although the role of positive symptom as a predictor of functional outcome is inconsistent in the literature (Green, 1996; Green et al., 2000). In addition, neurological examination abnormalities (NEA), which are subtle neurological abnormalities comprising impairments in motor function and sensory integration, and persistence of primitive reflexes, have been documented in first episode antipsychotic naïve patients (Sanders et al., 1994; Venkatasubramanian et al., 2003b) and patients with psychosis or BPD (Arabzadeh et al., 2014; Bora et al., 2018; Fountoulakis et al., 2018). NEA and the pathophysiology of schizophrenia may share a common genetic liability that affects the neurodevelopmental trajectory of this illness (Chan et al., 2009; Prasad et al., 2009).

Identifying individuals’ outcome trajectories at early stage of illness and potential risk factors associated with a poor outcome trajectory are of high priority because this knowledge can guide the development of individually tailored treatment and effective interventions that potentially alter the course of the disease. Many previous studies of functional outcome are limited by possible confounding factors, including prior use of neuroleptic medications (Arnold et al., 2015; Chakos et al., 1994; Keshavan et al., 1994) or illness chronicity (Schwarzkopf et al., 1990). Studies of first-episode, never treated patients can address these confounds. Also, most studies investigating functional outcomes start with clinical categorization and compare outcomes between different diagnostic groups, rather than relying on objective data driven approaches. Furthermore, when thinking about potential interventions, relatively few studies have considered potential socio-economic and personality characteristics between outcome trajectories. However, such analyses can help better understand the differences between trajectories and identify potential intervenable factors, such as substance use, that could be of interest to public policy.

Studies have applied unsupervised methods as exploratory tools to identify trajectory classes within the data. Statistical methods used to determine homogeneous group trajectories can be separated into two families. The first comprises model-based methods. These are related to mixture modelling techniques or latent class analysis, such as Group-Based Trajectory Modeling (Nagin and Odgers, 2010), growth mixture modeling (GMM) (Curran et al., 2010; Muth’en, 2004). Three recent reports have used GMM analysis examining patterns and predictors of trajectories for social and occupational functioning in FEP patients. Hodgekins et al. (2015) examined social recovery profile of FEP patients over 1-year period and found that male gender, younger age at onset of psychosis, more severe negative symptoms at baseline and poorer premorbid adjustment predicted poor recovery trajectory (Hodgekins et al., 2015). Velthorst et al. (2016) investigated long-term 20-year trajectories of social impairment in first-admission psychosis patients and identified four distinct but stable trajectories of social functional outcomes (Velthorst et al., 2016). Chang et al. (2018) investigated the social-occupational functional trajectories in Chinese FEP cohort over 3 years. The authors also found four distinct functional trajectories and that male gender, lower educational attainment, diagnosis of schizophrenia-spectrum disorders, as well as hospitalization upon first psychosis breakdown were associated with poor functional recovery. One essential step of growth models is the identification of the optimal functional form of the trajectory over time using the mean and covariance structure of the data, that is, the shape of the latent trajectories is specified in advance to fit a particular functional form of growth and formal tests are used to evaluate the fitness of the hypothesized model. As such, growth modeling approaches try to model trajectories and the group trajectories being theoretical constructs defined to take into account the heterogeneity in the data.

The second group of approaches relates to machine learning partitional classification analysis, such as K means clustering (Genolini and Falissard, 2010). The k-means algorithm uses variance distances or dissimilarity metrics such as the Euclidean distance metric between curves to identify and classify trajectories. The k-means algorithm corresponds to a non-parametric classification method: it searches the classification of the data that minimizes a specific within-group distance metric. The K means cluster modelling for longitudinal data (KmL) is thus an alternative classification to mixture modeling (Genolini and Falissard, 2011; Genolini et al., 2013; Subtil et al., 2017). Some potential advantages of k-means method are that: it is a non-parametric hill-climbing algorithm, does not impose assumptions regarding normality or the parameterization within the clusters or the shape of the trajectories, is likely to be more robust as regards numerical convergence, and is robust against mis-specification of model. The ability to classify outcome categories at an individual level is also likely to facilitate personalized approach to prediction and treatment. Although KmL approach has been used in biomedical studies to define clusters of patients with homogeneous trajectories of change (Pingault et al., 2013), or treatment response over time (Honer et al., 2015), to our knowledge, no report has applied objective data-driven KmL approach to identify homogeneous trajectories of functional outcome in patients with first episode psychosis (minimally treated with antipsychotics at study entry). Therefore, the primary aim of the study was to use unsupervised machine learning KmL method to empirically derive functional outcome trajectories of three core phenotypes (positive symptoms, negative symptoms, and NEA) in primarily unmedicated FEP patient cohort. Since the task was to cluster data on which no *a priori* hypothesis is available, and consequently would prefer to avoid the issues related to model selection using model-based procedures, the use of K means clustering is appropriate. The second aim was to characterize the socio-economic, personality, executive function, and clinical profiles of patients in each trajectory, in order to identify potentially modifiable processes or risk factors in FEP correlated with these different trajectories.

## Methods

### Subjects

Data were collected between 1996 and 2004 from The Pittsburgh First Episode of Psychosis (FEP) Longitudinal Cohort Study) (Keshavan et al., 1998). Patients were recruited from the inpatient and outpatient services of the Western Psychiatric Institute and Clinic, Pittsburgh. Patients with previously untreated psychosis were evaluated by trained clinicians using the Structured Clinical Interview for DSM-IV (SCID). Patients were included in the study if they were aged 15-45, had an IQ>75, met the DSM-IV criteria for a psychotic disorder, and had no or minimal prior treatment with neuroleptics, no significant medical or neurological illness, no history of head injury with loss of consciousness temporally related to psychosis onset, and no current substance abuse or dependence. DSM-IV diagnoses were derived in consensus diagnostic evaluations by the raters and senior MD or PhD level clinicians on the basis of all available clinical data and the SCID-P interview data. A total of 369 first episode patients (93% antipsychotic naive) with schizophrenia (SZ) (n= 221) or non-SZ psychoses (n= 144) were included in the baseline assessments. At study entry, patients who had received more than 2 weeks (in their lifetimes) of antipsychotic treatment were excluded. Patients were followed up at 4-8 weeks, 6-months, and one-year time point and treated with standard antipsychotic medications and supportive individual and/ or group psychotherapy in community treatment programs. The design, recruitment process and samples have been described in detail elsewhere (Keshavan et al., 2003; Li et al., 2011; Prasad et al., 2005). Patient social-demographic characteristics, clinical evaluation, and phenotypic assessments (described below) were recorded at each point.

### Clinical and phenotype assessments

Duration of Untreated Illness (DUI) was assessed in each case by the same raters using all clinical information, including medical records, reports by family members or significant others, and SCID interviews. The most likely date of illness onset, as defined by the onset of prodromal positive or negative symptoms, was then determined in consensus diagnostic conferences.

Psychopathological ratings were carried out by trained raters who also participated in the diagnostic meetings described above. Illness severity was assessed using the Global Assessment of Function (GAF) scale (Aas, 2010). GAF scores at 4-8 weeks, following stabilization of acute psychotic symptoms, were taken to reflect baseline level of functioning. Positive and negative symptoms were measured, respectively, using the Scale for the Assessment of Positive Symptoms (SAPS) and the Scale for the Assessment of Negative Symptoms (SANS) (Andreasen et al., 1990). General psychopathology symptoms were rated using the affective items from the Brief Psychiatric Rating Scale (Overall and Gorham, 1962) and the 24-item Hamilton Rating Scale for Depression (Hamilton, 1960). Scales were scored by averaging across items to produce an average rating for each of the symptom domains assessed. Ratings were conducted by trained and reliable (Kappa >.5) raters. Premorbid academic and social functioning were assessed using the Premorbid Adjustment Scale (PAS: (Cannon-Spoor et al., 1982) and were examined from childhood through late adolescence (Allen et al., 2001; Cannon et al., 1997).

Neurological examination abnormalities (NEA), also called neurological “soft” signs (NSS), were evaluated using the modified Buchanan–Heinrichs scale (Buchanan and Heinrichs, 1989; Sanders et al., 2006). NSS abnormality is predictive of neuropsychological performance in first episode psychosis (Arabzadeh et al., 2014; Mohr et al., 2003) and in chronic schizophrenic patients (Sewell et al., 2010), and is correlated with treatment response and recovery outcome (Prikryl et al., 2007). Our previous principal components analyses (Sanders et al., 2000; Venkatasubramanian et al., 2003b) revealed two dominant factors: repetitive motor sequencing (including fist-ring, alternating fist-palm, and rapid alternating movements) and cognitively demanding perceptual tasks (dominated by sensory/ audiovisual integration). The cognitive/perceptual neurological abnormalities factor correlates with neuroanatomical alterations and with neuropsychological assessments that tap into executive function and memory (Arabzadeh et al., 2014; Venkatasubramanian et al., 2003a). Studies also found a significant link from NSS to specific neurocognitive deficits (e.g., executive attention, verbal memory, and visual memory) and that more evidence of NSS is associated with more severe impairments in executive and memory functions, which are significant predators of functional outcomes(Arango et al., 1999). Given that functional outcome is of primary interest to the present study and given the evidence of significant relationships between NSS and executive function as well as between cognitive function and functional outcomes, we investigated the cognitive/perceptual neurological score in the analyses.

Personality assessments were carried out by a trained clinician (Keshavan et al., 2005). The Personality Disorder Evaluation, a semi-structured instrument (Loranger et al., 1991), was used and grouped personality dimensions into three main clusters, namely Cluster A (Paranoid, Schizoid, Schizotypal), Cluster B (Antisocial, Borderline, Histrionic, Narcissistic), and Cluster C (Avoidant, Dependent, Obsessive-compulsive) (Lenzenweger, 1999; Lenzenweger et al., 2007; Lenzenweger et al., 1997). The assessments were typically carried out before the consensus diagnostic conferences were held; the clinician was therefore unaware of the consensus diagnoses at the time of the personality assessment.

Executive functions were evaluated using the Wisconsin Card Sorting Test by a trained psychologist (WCST, Computerized Version 4.0)). Multiple categories were collected in this task but they are highly correlated. We selected number of perseverative errors as the main measure of executive function. Estimated premorbid IQ was assessed using the Ammon’s quick IQ test(Ammons and Ammons, 1962)

### Statistical analysis

#### Cluster Trajectory Analysis

K means cluster modelling for longitudinal data (K-means longitudinal 3D (kml3d), <http://CRAN.R-project.org/package=kml3d>), was used to identify distinct, homogeneous clusters of functional outcome trajectories over 4 assessment time points (baseline, 4-8 weeks, 6 month, 12 month). Because this study included three independent predictors (SAPS, SANS, NEA), three sets of two variable pairs (SAPS-GAF, SANS-GAF, NEA-GAF) k means clustering were performed in kml3d over 4 time points, enabling to study the joint evolution or complex interactions between variables over time (Genolini et al., 2015; Genolini et al., 2013). In addition, kml3d provides tools to visualize 3D dynamic graphs which can be exported in a 3D dynamic PDF. This visualization enables a better representation of the interaction between the two variable-trajectories. KmL provides substantially greater predictive power than single variable-trajectory (Shah et al., 2012) and is as efficient as the existing parametric algorithm on polynomial data, and is potentially more efficient on non-polynomial data (Genolini & Falissard, 2010). To evaluate the optimal number of cluster trajectories, we tested a number of models with different number of cluster trajectories (see supplementary material). Each model was repeatedly fitted with the number of clusters increasing step-wise from 2 to 6 using maximum likelihood criterion, computed using the KmL3D algorithm. We assessed model fit by using i) the Sum of Squared Error (SSE) within each cluster, ii) SSE between clusters, and iii) the Partitioning Around Mediods (PAM) method to measure how well individuals belong to a cluster trajectory and by using scree plots to visualize an “elbow,” or point representing the optimal number of clusters.

Patients with at least 3 assessment time points were retained in the analyses (N=129). The SAPS, SANS, and NEA cognitive/perceptual neurological score were used as the main predictors against the GAF score as the functioning outcome measure over 4 assessment time points. kml3d provides tools to visualize 3D dynamic graphs Missing values were imputed by carrying the last observation forward. Each of the three predictors (SANS, SAPS, NEA) was jointed with functional outcome variable GAF to derive functional trajectories in three separate analyses.

Once patients were assigned to specific trajectory groups, baseline information including demographics (sex, race, SES, IQ), executive function (WCST Number of perseverative errors), personality, and clinical features (diagnosis, histories of substance use, age of onset, premorbid function, duration of prodromal symptoms, brief psychotic rating Score, Hamilton rating scale for depression) were used to characterize each trajectory profile and to identify potential risk or protective factors for poor and good functional trajectories, respectively. ANOVA, t-test and χ^2^ tests were used to compare between and within trajectory groups.

## Results

A total of 369 patients were assessed in the baseline. Among them, 129 patients (SZ n= 82 & non-SZ psychoses n= 47) with at least 3 assessment data points were included in the kml3d analyses. Characteristics of the baseline sample and those who were included in the kml3d analyses were presented in Table 1. There were no significant differences between the two groups (Table 1).

**Table 1.**
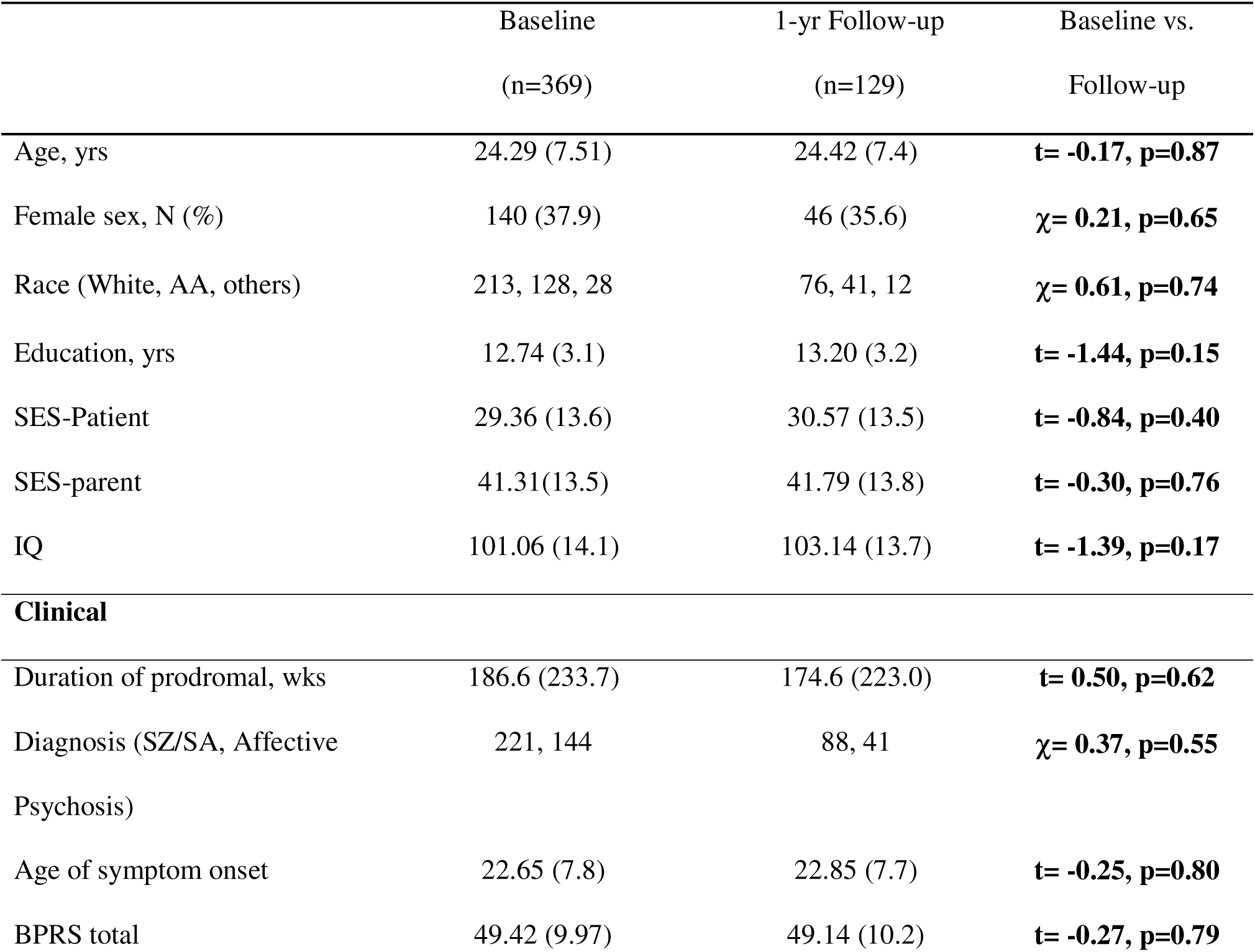
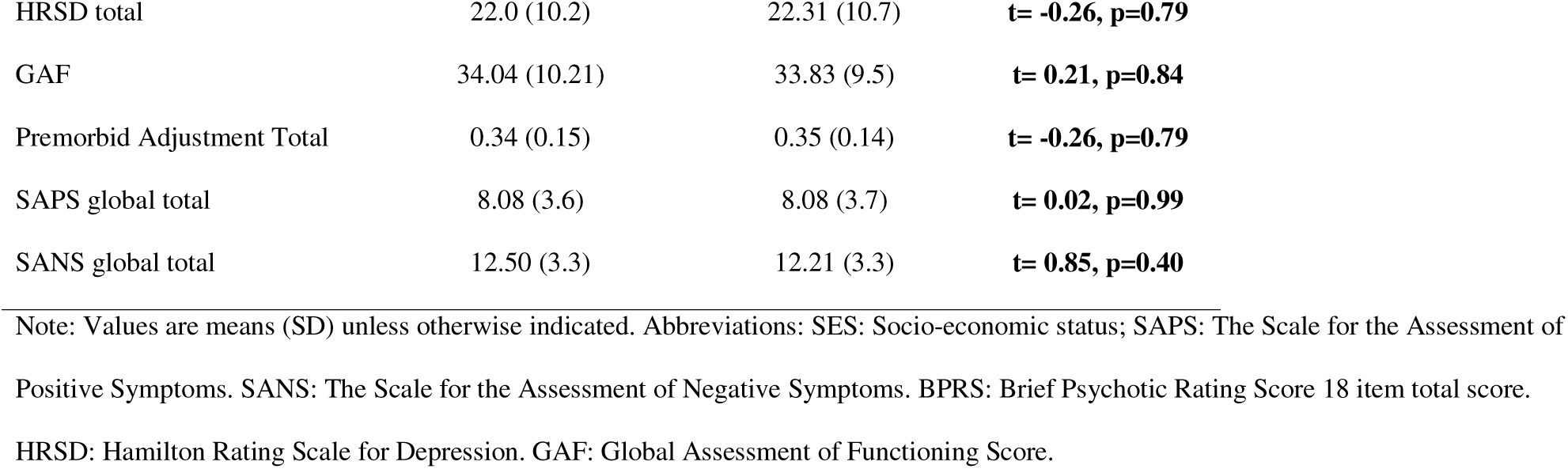
Socio-demographic and clinical Characteristics of Subject Groups

### Trajectory Identification

The four-trajectory solution was chosen as the best fit as this solution was the most consistent and captured cohesiveness and separation better than others (see supplementary material for detail and Figure S1 & S2). We labelled these four distinct functioning trajectories as *“Poor Outcome”, “Middle Outcome”, “Good Outcome”* and *“Catch-up*”. Patients in the good-outcome trajectory exhibited mild symptoms at baseline and the highest functioning a year later, whereas those in the poor-outcome trajectory had the most severe symptoms at baseline and recovered poorly a year later. Patients in the catch-up trajectory exhibited severe symptoms at baseline, indistinguishable to the poor-outcome patients, but were able to catch up in recovery to achieve good functioning a year later. Finally, patients in the middle-outcome trajectory showed moderate symptoms at baseline and partial functioning recovery a year later. The demographic and clinical characteristics of each trajectory and statistical comparisons between Good and Poor outcomes are presented in Table 2-4. Comparisons among other groups are presented in the Supplementary Materials. We provide an online file which allows the reader to explore the three-dimensional trajectories in a dynamic fashion (<kml3d.saps.html>kml3d.sans.html><kml3d.nes.html>).

**Table 2.** The four-trajectory model using SANS symptoms as predictor against GAF

|  | Poor (A)<br>N=48 | Middle (B)<br>N=32 | Catch-up (C)<br>N=27 | Good (D)<br>N=22 | Group<br>Comparison | Good vs. Poor |
| --- | --- | --- | --- | --- | --- | --- |
| SCZ (%) | 81% | 37% | 70% | 55% | $\chi^2 = 15.58$;<br>P= 0.001 | $\chi^2 = 5.44$;<br>P= 0.020 |
| Affective psychosis (%) | 19% | 62% | 30% | 45% |  |  |
| SEX Male (%) | 77% | 56% | 55% | 59% | $\chi^2 = 4.68$;<br>P=0.2 | $\chi^2 = 2.39$;<br>P= 0.12 |
| White | 48% | 59% | 74% | 64% | $\chi^2 = 15.64$;<br>P=0.016 | $\chi^2 = 7.62$;<br>P= 0.020 |
| AA | 48% | 31% | 15% | 18% |  |  |
| Others | 2% | 10% | 11% | 18% |  |  |
| Avg IQ score | 98.68 (13) | 104.76 (15) | 105.19 (13) | 109.06 (12) | F=3.19;<br>P=0.02 | t = 2.83;<br>P=0.006* |
| Patient SES | 27 | 32 | 37 | 28.5 | F=2.69;<br>P=0.049 | t = 0.23;<br>P=0.81 |
| Parent SES | 40 | 41 | 42.5 | 44 | F=0.36;<br>P=0.78 | t = 1.04;<br>P=0.30 |
| Age onset | 22.12 (6.9) | 22.61 (7.2) | 26.39 (9.4) | 19.16 (7.4) | F=4.21;<br>P=0.007 | t = -1.71;<br>P=0.09 |
| Substance use Frequency | 49% | 42% | 28.5% | 23.8% | $\chi^2 = 5.49$;<br>P=0.14 | $\chi^2 = 3.84$;<br>P= 0.05 |
| BPRS-18 items | 52.59 (9.8) | 46.48 (10.2) | 50.62 (8.8) | 43.19 (9.7) | F=5.66;<br>P=0.001 | t = -3.67;<br>P=0.0005* |
| HRSM-24 items | 22.36 (10.1) | 23.5 (12.0) | 24.44 (10.1) | 16.81 (8.8) | F=1.92;<br>P=0.13 | t = -1.87;<br>P=0.07 |
| WCST-perseverative errors | 25.15 (12.3) | 19.27 (13.6) | 20.22 (13.3) | 22.12 (23.0) | F=0.64;<br>P=0.59 | t = -0.45;<br>P=0.65 |
| PMAS-CHILD | 0.30 | 0.30 | 0.33 | 0.28 | F=0.09;<br>P=0.96 | t = -0.24;<br>P=0.81 |
| PMAS-TOTAL | 0.38 | 0.32 | 0.35 | 0.32 | F=1.10;<br>P=0.35 | t = -1.61;<br>P=0.12 |
| Duration of Prodromal (wk) | 148.20 (152.9) | 215.42 (276.3) | 147.95 (220.4) | 210.06 (274.8) | F=0.67;<br>P=0.57 | t = 1.13;<br>P=0.26 |
| CLUSTER-A | 16.9 (11.2) | 14.2 (10.1) | 9.5 (6.3) | 9.6 (7.5) | F=2.15;<br>P=0.10 | t = -1.99;<br>P=0.056 |
| CLUSTER-B | 5.65 (6.7) | 15.35 (17.2) | 5.22 (6.3) | 7.08 (6.9) | F=3.29;<br>P=0.026 | t = 0.55;<br>P=0.59 |
| CLUSTER-C | 9.00 (6.3) | 8.94 (6.2) | 6.89 (5.5) | 5.93 (4.5) | F=1.67;<br>P=0.18 | t = -2.11;<br>P=0.04 |

### SANS as Predictor of Functioning Trajectories

Patients in the good-outcome trajectory (Table 2, Figure 1, Purple color [D]) had the highest premorbid IQ score and parental SES but the lowest age of psychotic symptom onset, BPRS and HRSM scores, premorbid childhood abnormalities and personality cluster A traits. They were the least likely to use substances than patients in the other trajectories. Patients in the Catch-up trajectory (Figure 1, Cyan color [C]) were predominately diagnosed with SZ and were more likely to be Caucasian. They share similar characteristics with good-outcome patients in terms of SES, IQ, low personality cluster A traits, and low substance use, but differed in terms of psychotic symptom age onset, HRSM scores, and childhood premorbid abnormalities.

**Figure 1.**
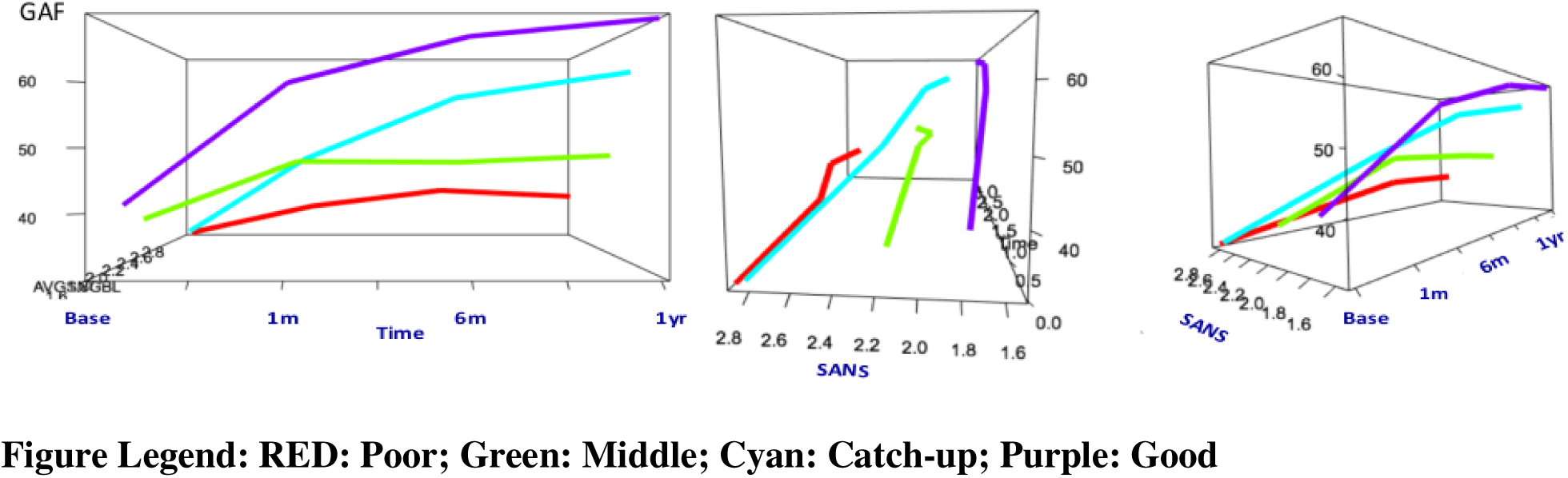
SANS symptoms as predictor against GAF.

Patients in the middle-outcome trajectory (Figure 1, Green [B]) were more likely to have affective psychosis (62%) diagnosis. These patients performed well (fewest errors) on WCST. Patients in the poor-outcome trajectory (Figure 1, Red [A]) were primarily diagnosed with SZ and more likely to be male. These patients had the lowest IQ score and came from lower SES strata. In addition to severe negative symptoms, they also exhibited high premorbid abnormalities, high personality cluster A and C traits, performed the worst in WCST, and used substance the most than patients in the other trajectories (also see <kml3d.sans.html>).

ANOVA results showed significant group differences in the distribution of diagnoses, race, IQ, BPRS score, substance use history, and personality trait. Patients in the good outcome trajectory had more evenly distributed diagnosis, more Caucasian, higher IQ, lower psychotic (BPRS) symptoms, and lower Cluster B personality trait (Antisocial, Borderline, Histrionic, Narcissistic) than those in the poor trajectory group (Table 2).

### SAPS as Predictor of Functioning Trajectories

Patients in the good-outcome trajectory (Table 3, Figure 2, Green color [B]) were more likely to be Caucasian (66%), had higher IQ score, parental SES, and premorbid function and lower BPRS and HRSM scores, personality cluster A traits, and used substance less than patients in the other trajectories. Patients in the Catch-up Trajectory (Figure 2, Cyan color [C]) were predominately diagnosed with SZ. They had higher personal SES and HRSM score, older psychotic symptom onset, and lower personality cluster B traits than patients in the other trajectories. Patients in the poor-outcome trajectory (Figure 2, Purple color [D]) were primarily diagnosed with SZ (86%), more likely to be male (81%) or African American (57%), a lower IQ score and came from lower SES strata. They also exhibited high personality cluster B traits, performed the worst in WCST task, and used substances the most than patients in the other trajectories.

**Table 3.**
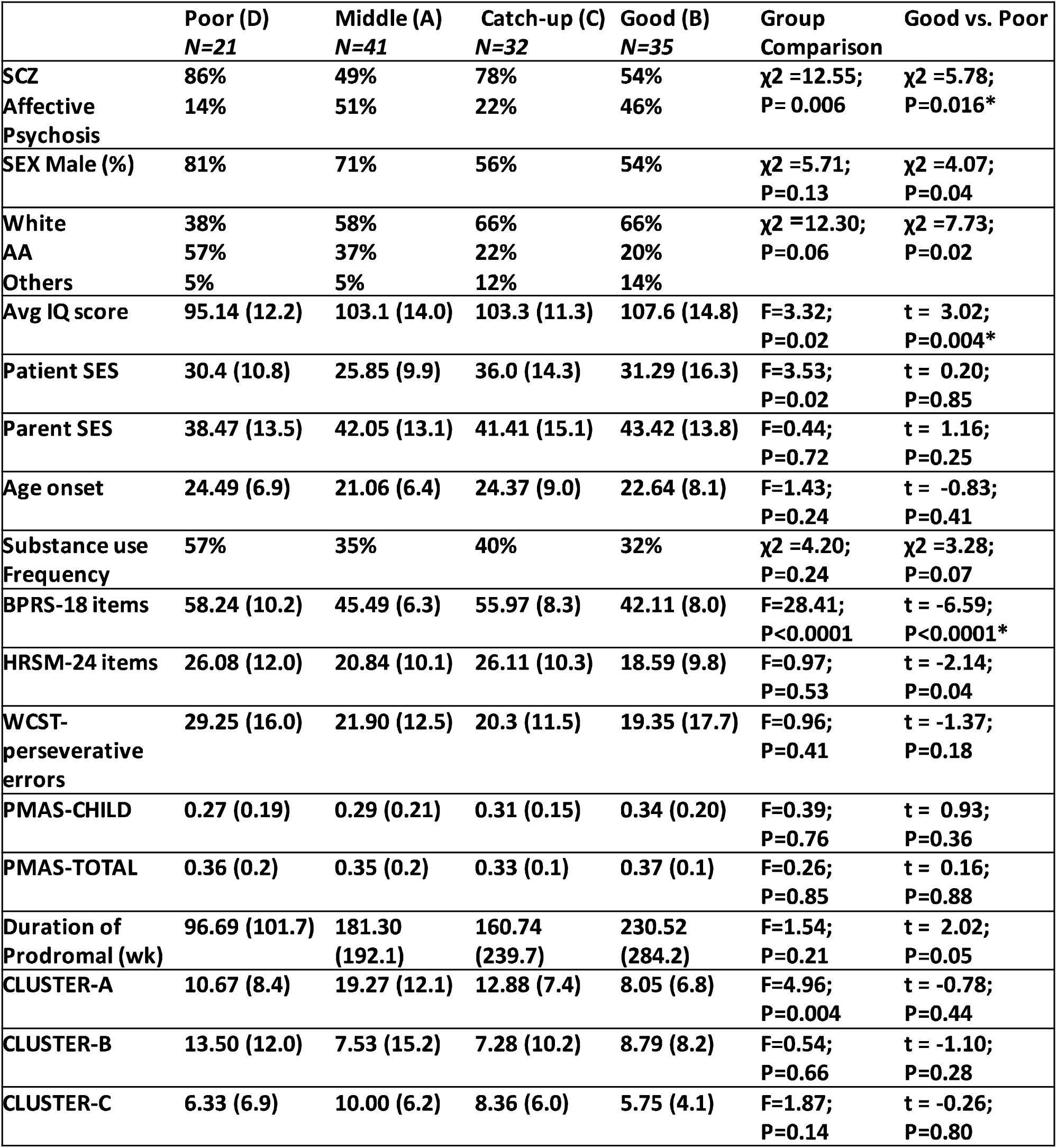
The four-trajectory model using SAPS symptoms as predictor against GAF

**Figure 2.**
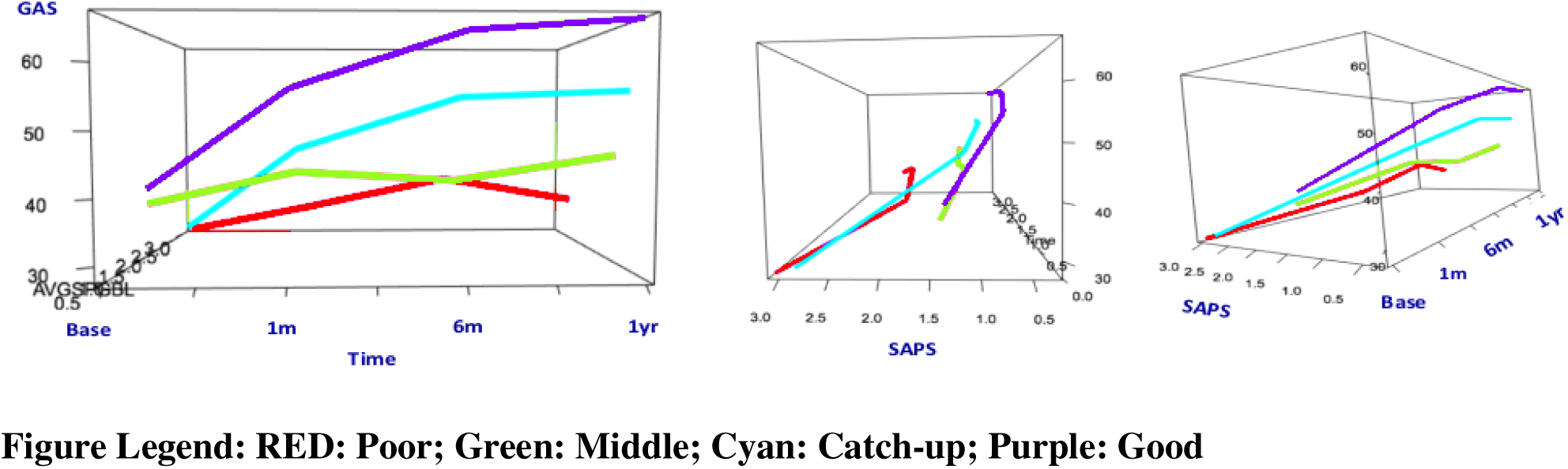
SAPS symptoms as predictor against GAF.

Significant group differences were fund between good and poor outcome trajectory in the distribution of diagnoses, IQ, patent SES, BPRS, and personality cluster A traits. Patients in the good outcome trajectory had more evenly distributed diagnosis, were more often Caucasian had higher IQ, lower psychotic (BPRS), and lower depression and anxiety (HRSM) symptoms than those in the poor trajectory group (Table 3).

### NEA (NSS) as Predictor of Functioning Trajectory

Consistent with results obtained from SANS and SAPS, good-outcome patients (Figure 3, Red color [A]) were more likely to be Caucasian, had higher IQ score and SES, performed well on WCST, and used substances the least than patients in the other trajectories. Patients in the catch-up trajectory (Figure 3, Cyan color [C]) were predominately diagnosed with SZ and were more likely to be Caucasian, had higher premorbid function, lower age onset, lower personality A and C traits than patients in the other trajectories. Patients in the poor-outcome trajectory (Figure 3, Purple [D]) were primarily diagnosed with SZ (86%), more likely to be male (74%), had lower IQ, performed poorly on WCST, came from lower SES strata, and were used substance the most than patients in the other trajectories.

**Figure 3.**
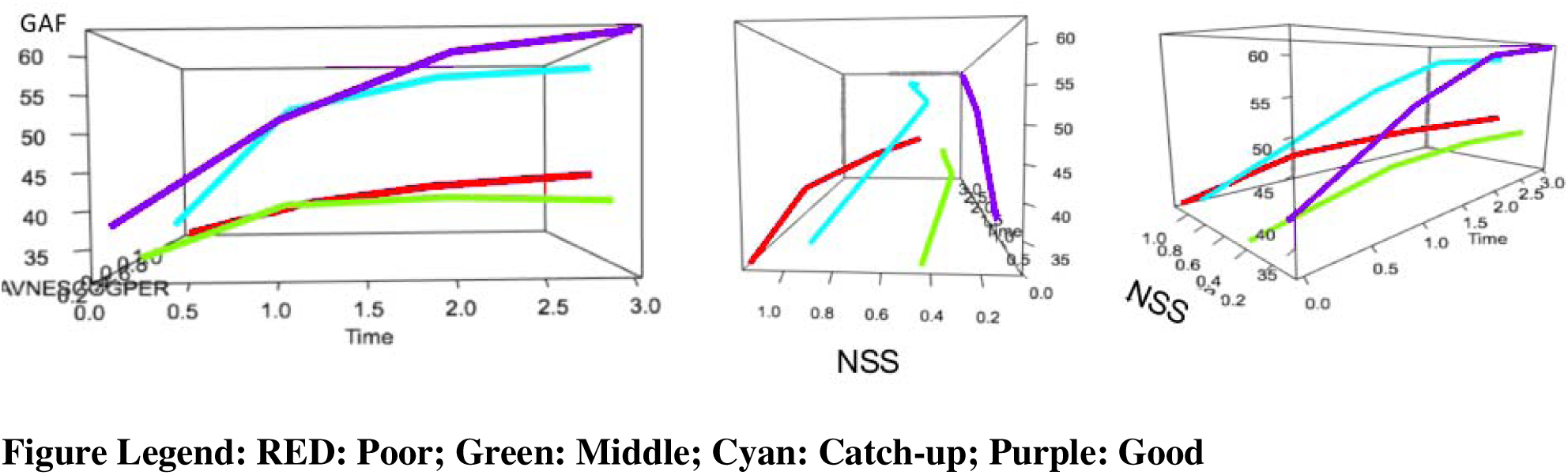
NSS symptoms as predictor against GAF.

Significant group differences were found in the distribution of diagnoses, IQ, parent SES, and BPRS. The good outcome trajectory had more evenly distributed diagnosis, higher IQ, and lower psychotic (BPRS) than those in the poor trajectory group (Table 4).

**Table 4.** The four-trajectory model using NSS at baseline as predictor against GAF

|  | Poor (D)<br>N=21 | Middle (B)<br>N=32 | Catch-up (C)<br>N=23 | Good (A)<br>N=44 | Group<br>Comparison | Good vs. Poor |
| --- | --- | --- | --- | --- | --- | --- |
| SCZ | 86% | 63% | 70% | 48% | $\chi^2 = 9.42$;<br>P= 0.02 | $\chi^2 = 8.55$;<br>P= 0.003* |
| BPD | 14% | 37% | 30% | 52% |  |  |
| SEX Male (%) | 74% | 58% | 67% | 61% | $\chi^2 = 2.09$;<br>P=0.56 | $\chi^2 = 1.82$;<br>P= 0.18 |
| White | 52% | 50% | 71% | 69% | $\chi^2 = 20.84$;<br>P=0.05 | $\chi^2 = 4.91$;<br>P= 0.09 |
| AA | 43% | 47% | 17% | 17% |  |  |
| Others | 5% | 3% | 12% | 14% |  |  |
| Avg IQ score | 89.62 (11.07) | 103.40 (11.9) | 100.28 (11.0) | 111.92 (12.7) | F=15.37;<br>P<0.0001 | t = 6.43;<br>P<0.0001* |
| Patient SES | 27.2 (8.9) | 29.5 (9.6) | 28.4 (15.6) | 34.4 (16.2) | F=1.77; P=0.16 | t = 1.82;<br>P=0.07 |
| Parent SES | 33.3 (12.4) | 41.0 (13.5) | 35.8 (12.2) | 47.4 (12.6) | F=6.09;<br>P=0.0008 | t = 3.63;<br>P=0.0006* |
| Age onset | 25.2 (9.6) | 21.6 (5.5) | 21.0 (8.3) | 23.0 (7.7) | F=1.34; P=0.27 | t = -0.99;<br>P=0.33 |
| Substance use | 48% | 44% | 35% | 30% | F=2.71; P=0.44 | $\chi^2 = 2.03$;<br>P= 0.15 |
| BPRS-18 items | 52.90 (11.2) | 51.03 (8.0) | 48.0 (11.9) | 45.38 (8.7) | F=3.61; P=0.01 | t = -2.92;<br>P=0.005* |
| HRSM-24 items | 24.0 (10.1) | 25.71 (12.6) | 20.0 (10.2) | 20.63 (9.1) | F=1.57; P=0.20 | t = -1.13;<br>P=0.26 |
| WCST-perseverative<br>errors | 32.25 (17.6) | 22.89 (11.3) | 24.5 (17.7) | 14.79 (11.6) | F=2.49; P=0.07 | t = -2.67;<br>P=0.01 |
| PMASCHILD | 0.36 (0.2) | 0.28 (0.2) | 0.30 (0.2) | 0.30 (0.1) | F=0.47; P=0.70 | t = -0.82;<br>P=0.42 |
| PMAS TOT | 0.39 (0.2) | 0.35 (0.1) | 0.32 (0.1) | 0.34 (0.1) | F=0.55; P=0.65 | t = -0.99;<br>P=0.33 |
| Duration of<br>Prodromal (wk) | 124.07 (127.2) | 212.23 (212.8) | 173.97 (183) | 200.06(294.8) | F=0.72;<br>P=0.54 | t = 1.12;<br>P=0.26 |
| CLUSTER-A | 15.7 (11.1) | 17.07 (11.4) | 9.0 (6.8) | 11.31 (8.2) | F=2.15; P=0.10 | t = -1.30;<br>P=0.20 |
| CLUSTER-B | 4.4 (3.3) | 10.3 (12.1) | 11.07 (11.0) | 7.88 (12.8) | F=0.80; P=0.50 | t = 0.84;<br>P=0.41 |
| CLUSTER-C | 8.4 (6.0) | 9.86 (6.3) | 4.23 (3.2) | 8.58 (5.8) | F=2.63; P=0.06 | t = 0.08;<br>P=0.94 |

## Discussion

In this study, we employed machine learning unsupervised kml3d clustering methods to derive distinct, homogeneous clusters of short-term functional outcome trajectories in a sample of patients with FEP. These patients were antipsychotic-naïve or minimally antipsychotic-treated at study entry and were longitudinally followed at 4-8 weeks, 6 months and 12 months later. Three core phenotypes were used, each as an independent predictor in separate kml3d models, to examine the joint trajectories of each predictor with functional outcome. To preserve a larger sample size and minimize the influence of imputation on the results (i.e., more accurate standard error estimates) (Twisk and de Vente, 2002), we included only patients with at least 3 assessment time points rather than imputing all the missing data in the kml3d analyses. Although there have been many studies of outcome in SZ (Carpenter and Kirkpatrick, 1988; Levine et al., 2011), few have used a data driven approach to empirically detect distinct outcome trajectories while capturing the longitudinal course of functional levels across the entire study period. Trajectory analysis is a useful strategy to reduce heterogeneity and provide insight into clinically meaningful subgroups of patients (Honer et al., 2015).

Results indicated that a four-cluster trajectory was the best fit. Importantly, the four trajectories identified in each predictor model are highly consistent and concordant in terms of the overall profiles (trajectory shapes), patterns, and the sample characteristics. We labelled the 4 distinct functional recovery trajectories as: (1) Good Outcome--individuals had the mildest symptoms or neurological abnormality at study entry and recovered well to achieve the highest functioning a year later; (2) Poor Outcome--individuals had the most severe symptoms or neurological abnormality at study entry and recovered poorly to achieve the lowest functioning a year later; (3) Intermediate Outcome -- individuals had moderate symptoms or neurological abnormality at study entry and recovered to achieve intermediate level of functioning a year later; (4) Catch-up--individuals had severe symptoms or neurological abnormality at study entry but were able to “catch-up” in recovery a year later to achieve global functioning level similar to individuals in the good outcome trajectory.

The Catch-up Outcome trajectory is the most intriguing. At the study entry, individuals in the Catch-up trajectory were indistinguishable from those in the Poor Outcome trajectory, exhibiting severe positive and negative symptoms and cognitive/perceptual neurological abnormality. However, the recovery path was very different. Patients in the Catch-up trajectory were able to recover quickly and to achieve functioning level akin to those in the good-outcome a year later (at 12-month follow-up Poor: GAF score between 38.8 and 45.5; Catch-up: GAF score between 56.3 and 62.3). In addition, the Catch-up trajectory consistently emerged in each predictor model. The characteristics of individuals in the Catch-up trajectory appear to be more female, Caucasian, having higher IQ, higher executive function, having better premorbid adjustment, coming from higher socio-economic background, as well as having less substance use history as compared with individuals in the Poor Outcome trajectory. These differences in characteristics between Poor and Catch-up were robustly consistent across all three predictor models.

On the other hand, across all three models patients in the Good Outcome and Catch-up appear to share similar socio-economic and cognitive function profile, such that patients with good outcome were also more likely to be female, Caucasian, having higher IQ and higher executive function, having lower psychopathology and personality cluster A (Paranoid, Schizoid, Schizotypal) traits, coming from higher socio-economic background, as well as having less substance use history than those in the Poor trajectory.

In our study, the same diagnoses did not have a strong predictive value for the different trajectories. A majority of individuals in the Poor Outcome and Catch-up trajectories have diagnosis of SZ, whereas patients in the Good Outcome and Middle trajectories include both SZ and affective psychosis. Biologically based phenotypes at baseline may allow better predictions (Clementz et al., 2016; Hall et al., 2012). Many clinical and neurobiological alterations collectively contribute to variance in functioning (Allott et al., 2011). Our results showed that multivariate models that integrate diverse domains of risk predictors at baseline can identify neurobiologically homogeneous sub-groups, which provide substantially greater predictive power for functional trajectory.

Our results suggest that being male, being an ethnic minority, coming from disadvantaged socio-economic strata, having a poor baseline cognitive function, high in mood (depression/anxiety) symptoms, high in specific personality trait (particularly paranoid/schizotypal), and heavy substance use may be risk factors associated with a poorer functioning recovery. These findings are consistent with previous literature which identifies gender (Cotton et al., 2009); ethnic minority status (Hodgekins et al., 2015; Li et al., 2011; Morgan et al., 2008); premorbid adjustment (Addington and Addington, 2005); and personality (Compton et al., 2015; Lysaker et al., 1998; Lysaker and Davis, 2004) as predictors of poor recovery following psychosis. Our results also indicate that substance use history contributes significantly in predicting functioning recovery.

The identification of four-trajectory solution as the best model accords with both Velthorst et al., 2016 and (Chang et al., 2018), although different analysis approaches were used. Velthorst and colleagues detected four stable trajectories of preserved, moderately, severely, and profoundly impaired social functioning across diagnoses over 20 years follow-up (Velthorst et al., 2016). Given the short-term follow up period in the present study, we were unable to directly compare with Velthorst in terms of long-term trajectory patterns. Chang and colleagues employed the growth mixture modeling (GMM), to explore social-occupational functional trajectories in patients with first-episode non-affective psychosis across 3-year follow-up (Chang et al., 2018). The overall patterns of the ‘gradually improved’, the ‘early improved’ and the ‘persistently poor’ trajectories reported by Chang and colleagues resemble the ‘catch-up’, ‘good outcome’, and ‘poor outcome’ trajectories identified in our study, respectively. However, the ‘improved-deteriorated’ trajectory identified in Chang et al study was not found in our study. Several reasons might, in part, account for the discrepancy. For example, our sample cohort included both affective and non-affective psychosis patients whereas Chang et al study included only non-affective psychosis patients. GAF score was used in our study whereas SOFAS score was used in Chang et al study. Another report by Hodgekins and colleagues focusing on social functioning recovery in individuals with FEP over a12-month period. These authors found three types of social recovery profile: Low Stable, Moderate-Increasing, and High-Decreasing (Hodgekins et al., 2015). The ‘low stable’ profile is the most consistent with the ‘poor-outcome’ trajectory identified in our study. The ‘moderate-Increasing’, and ‘high-decreasing’ profiles are different. The present study differs from Hodgekins et al report in many aspects, including functioning assessments (Time use survey vs. GAF) and primary focus in functioning (specific social functioning vs. global impression of functioning). Functional outcome represents a multifaceted variable relying on different constructs and depending on several clinical neuropsychological and environmental factors (Bechi et al., 2017). It is likely different patterns of recovery trajectory exist in individuals with FEP depending specific aspect of functional outcome and tools used to assess these functional outcomes.

Several explanations may account for the lack of functional improvement in patients with Poor Outcome trajectory. Patients in this group may have less access to resources, medication, and/or necessary treatment because of being in an unfavorable socio-economic circumstance. In addition, evidence suggests that in the African American population, psychiatric disorders including psychosis have greater stigma attached to them. Blacks with psychiatric disorders are more likely to be viewed as morally inferior which may result in a lessened tendency to adhere to treatment (Alvidrez, 1999). Financial burden, access difficulties, and stigma may reduce patients’ access to better treatment and increase their isolation and hopelessness, thereby decreasing or slowing down their recovery (Gary, 2005). Furthermore, personality variables have been linked with patterns of substance abuse (Lysaker and Davis, 2004; Van Os and Jones, 2001), which may impede treatment, worsen cognitive impairment, reduce motivation, and slow down recovery. Finally, it is possible that patients in the poor outcome trajectory may differ neurobiologically (and may represent distinct biotypes) from those in the good outcome categories. This possibility can be empirically tested by longitudinal prospective studies with neuroimaging and electrophysiological studies carried out at baseline.

Our results should be interpreted in light of some limitations. First, the current study focused on short-term functioning trajectory during the first 12 months of psychosis onset. Therefore, it is unclear what happens after the first year in terms of functional recovery. However, one study has shown that the early clinical course is a good predictor for the long-term course (Rund et al., 2016). Second, the differences between trajectories were examined using baseline characteristics only. Factors such as engagement with services, family or social support, treatment adherence, or medication use during the follow-up period were not directly assessed. Third, the present study enrolled 369 patients at baseline, only 129 patients’ data (35%) were used for further analysis. Although we did not see any clinical or demographic differences between those with vs without follow-up data, it is possible that this large attrition may impact generalizability of our findings. Finally, as in any exploratory modeling, identification of trajectory classes is affected by sample size. Future study with larger sample is warranted to validate the results.

In summary, we identified four distinct and consistent patterns of functional trajectories across three independent predictors. Individuals with male gender; ethnic minority status; low premorbid adjustment; low executive function/IQ, low SES, personality disorder, substance use history may be risk factors for poor recovery following psychosis. Our findings suggest that functional recovery trajectories appear to be complicated because many psychological and environmental changes occur during the first year after illness onset. Clearly, data driven individual based trajectory analysis can facilitate the understanding of the population heterogeneity and the identification of risk factors associated with poor recovery path. One logical and important next step is to be able to evaluate candidate models that can accurate predict individual level outcomes for improving the quality in personalized treatment and targeted intervention.

## Financial Disclosures

The authors reported no biomedical financial interests or potential conflicts of interest.

## Role of the Funding Source

This publication was supported by funds received from National Institute of Mental Health grants NIMH MH45156, MH64023 NIH/NCRR/GCRC grant #M01 RR00056 (MSK). and [R01MH109687]: Mei-Hua Hall, PI; [K24MH104449]: Dost Öngür, PI

## Conflict of interest

One of the authors is an editor of this journal. All editorial process for this paper is handled by another editor.

## Supporting information

Supplementary Material: Cluster Trajectory Evaluation

## Acknowledgments

We thank Drs. Nina Schooler, Cameron carter, Raymond Cho, MD, Gretchen Haas PhD, and the clinical core staff of the Center for the Neuroscience of Mental Disorders (David Lewis MD, Director). We thank Kevin Eklund and Elizabeth Radomsky for their assistance in diagnostic, psychopathological and neuropsychological assessments, and Jean Miewald for help in data management..

