## Supplementary Material: Cluster Trajectory Evaluation for "Longitudinal Trajectory of Early Functional Recovery in Patients with First Episode Psychosis"

To evaluate the optimal number of cluster trajectories, we assessed model fit by using i) the Sum of Squared Error (SSE) within each cluster, ii) SSE between clusters, and iii) the Partitioning Around Mediods (PAM) method at baseline to measure how well individuals belong to a cluster trajectory and by using scree plots to visualize an “elbow,” or point representing the optimal number of clusters. Each model was repeatedly fitted with the number of clusters increasing step-wise from 2 to 6 using maximum likelihood criterion, computed using the KmL3D algorithm.

*The Sum of Squared Error (SSE) within each cluster*

The SSE within each cluster measures how well individuals belong to a cluster trajectory (cohesiveness).   An obvious bend or “elbow” indicates the “optimal” number of clusters that fit the data. For both SANS-GAS and SAPS-GAS, the optimal number of clusters appears to be 4, where there is an “elbow” (Figure S1a & b).

*The Sum of Squared Error (SSE) between clusters*

The SSE between clusters ascertains how well the clusters are separated, and this number is to be maximized.  For SANS-GAS, the cluster SSE between increases with the number of clusters (Figure S1c).  For SAPS-GAS the optimal number of clusters appears to be 4 (Figure S1d).

Figure S1. Results of SSE within each cluster (Top) and SSE between clusters (Bottom) plots

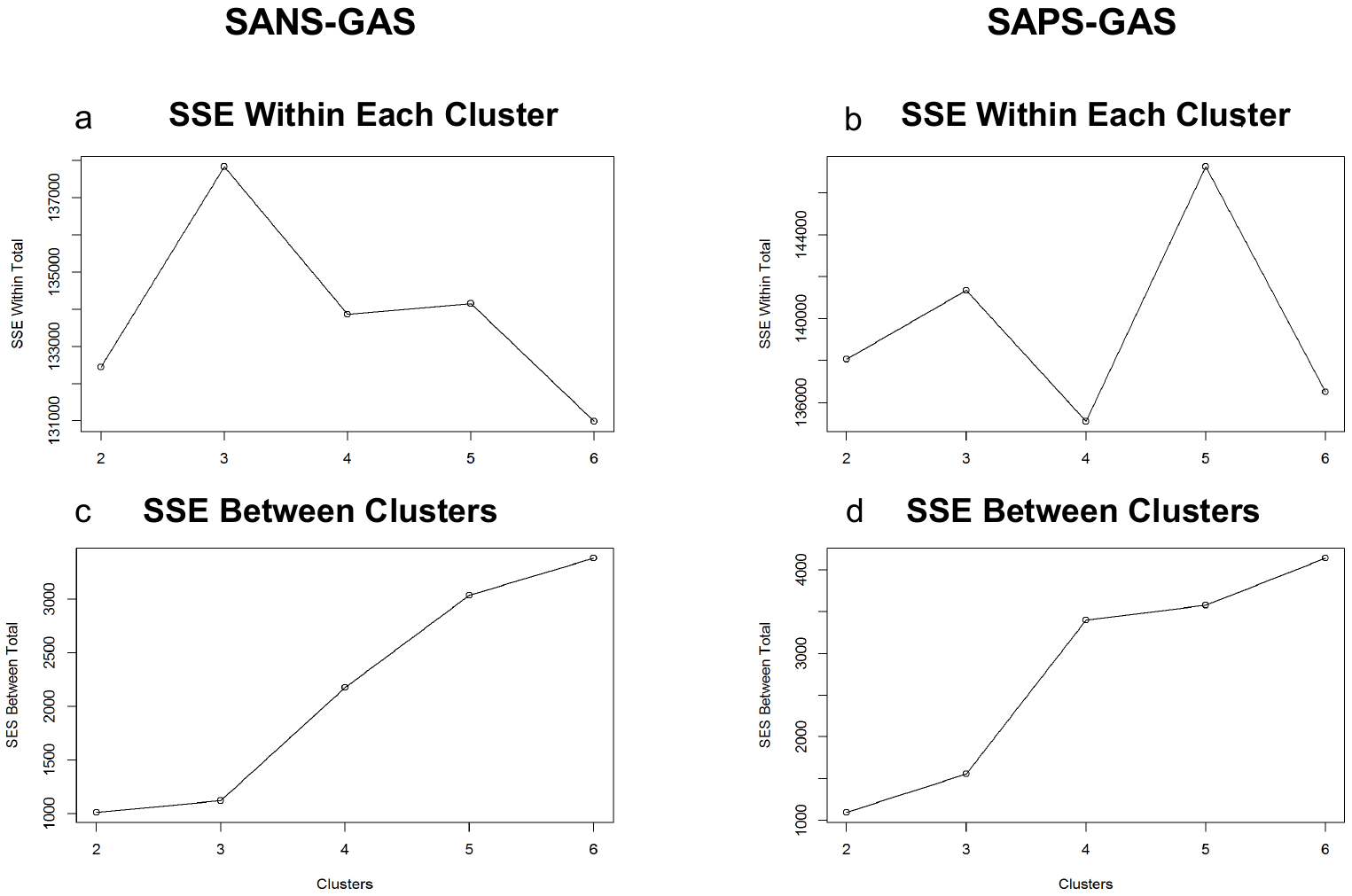

*Partitioning Around Mediods (PAM) clustering*

PAM clustering analysis chooses actual data points as medioid centers, and works with an arbitrary distance matrix (Kaufman and Rousseeuw, 1987).  PAM clustering for SANS-GAS at baseline indicates the optimal number of clusters is 4. Likewise, for SAPS-GAS, PAM calculates 4 clusters are the optimal number of clusters (Figure S2).

Figure S2. Results of PAM analysis plots

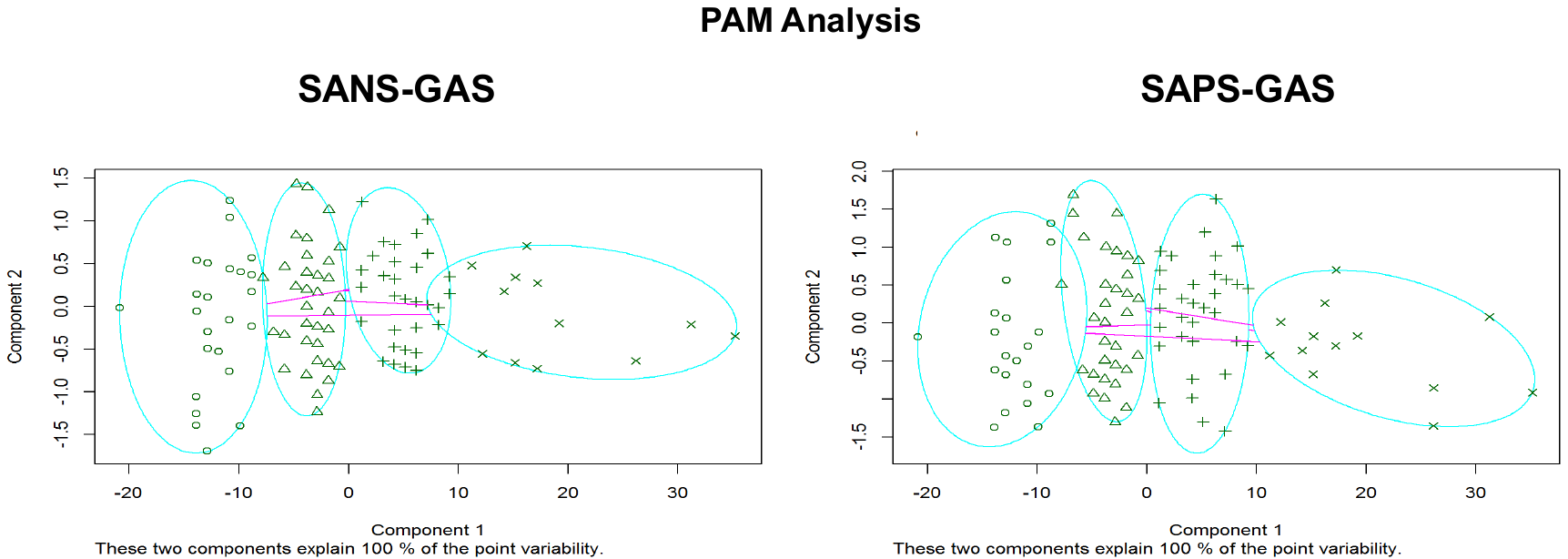

Taken together, these cluster evaluation methods indicate that, for both SANS-GAS and SAPS-GAS, the optimal number of clusters appears to be 4.

Table S1. SANS as predictor: Comparisons among trajectory groups

|  | Good vs. catch-up | Good vs. Middle | Catch-up vs. Middle | Catch-up vs. Poor | Middle vs. Poor |
| --- | --- | --- | --- | --- | --- |
| SCZ (%) | N.S. | N.S. | 2 =6.35;  P= 0.01 | N.S. | χ2 =13.73;  P< 0.001 |
| Affective psychosis (%) |  |  |  |  |  |
| SEX Male (%) | N.S. | N.S. | N.S. | χ2 =3.91;  P= 0.04 | N.S. |
| White | N.S. | N.S. | N.S. | χ2 =6.55;  P= 0.009 | N.S. |
| AA |  |  |  |  |  |
| Others |  |  |  |  |  |
| Avg IQ score | N.S. | N.S. | N.S. | N.S. | N.S. |
| Patient SES | N.S. | N.S. | N.S. | t = 3.04;  P=0.003 | N.S. |
| Parent SES | N.S. | N.S. | N.S. | N.S. | N.S. |
| Age onset | t = -3.09;  P=0.004 | N.S. | N.S. | N.S. | N.S. |
| Substance use Frequency | N.S. | N.S. | N.S. | N.S. | N.S. |
| BPRS-18 items | t = -2.75;  P=0.009 | N.S. | N.S. | N.S. | N.S. |
| HRSM-24 items | t = -2.47;  P=0.02 | N.S. | N.S. | N.S. | N.S. |
| WCST-perseverative errors | N.S. | N.S. | N.S. | N.S. | N.S. |
| PMAS-CHILD | N.S. | N.S. | N.S. | N.S. | N.S. |
| PMAS-TOTAL | N.S. | N.S. | N.S. | N.S. | N.S. |
| Duration of Prodromal (wk) | N.S. | N.S. | N.S. | N.S. | N.S. |
| CLUSTER-A | N.S. | N.S. | N.S. | N.S. | N.S. |
| CLUSTER-B | N.S. | N.S. | t = -2.29;  P=0.03 | N.S. | t = 2.12;  P=0.04 |
| CLUSTER-C | N.S. | N.S. | N.S. | N.S. | N.S. |

Table S2. SAPS as predictor: Comparisons among trajectory groups

|  | Good vs. catch-up | Good vs. Middle | Catch-up vs. Middle | Catch-up vs. Poor | Middle vs. Poor |
| --- | --- | --- | --- | --- | --- |
| SCZ (%) | N.S. | N.S. | χ2 =6.54;  P= 0.01 | N.S. | χ2 =7.98;  P= 0.005 |
| Affective psychosis (%) |  |  |  |  |  |
| SEX Male (%) | N.S. | N.S. | N.S. | N.S. | N.S. |
| White | N.S. | N.S. | N.S. | χ2 =5.97;  P= 0.02 | N.S. |
| AA |  |  |  |  |  |
| Others |  |  |  |  |  |
| Avg IQ score | N.S. | N.S. | N.S. | t = 2.38;  P=0.02 | t = 2.08;  P=0.04 |
| Patient SES | N.S. | N.S. | t = 3.52;  P=0.0008 | N.S. | N.S. |
| Parent SES | N.S. | N.S. | N.S. | N.S. | N.S. |
| Age onset | N.S. | N.S. | N.S. | N.S. | N.S. |
| Substance use Frequency | N.S. | N.S. | N.S. | N.S. | N.S. |
| BPRS-18 items | t = -6.84;  P<0.001 | N.S. | t = 6.06;  P<0.001 | N.S. | t = -6.10;  P<0.001 |
| HRSM-24 items | t = -2.82;  P=0.007 | N.S. | N.S. | N.S. | N.S. |
| WCST-perseverative errors | N.S. | N.S. | N.S. | N.S. | N.S. |
| PMAS-CHILD | N.S. | N.S. | N.S. | N.S. | N.S. |
| PMAS-TOTAL | N.S. | N.S. | N.S. | N.S. | N.S. |
| Duration of Prodromal (wk) | N.S. | N.S. | N.S. | N.S. | N.S. |
| CLUSTER-A | N.S. | t = -3.49;  P=0.001 | N.S. | N.S. | N.S. |
| CLUSTER-B | N.S. | N.S. | N.S. | N.S. | N.S. |
| CLUSTER-C | N.S. | N.S. | N.S. | N.S. | N.S. |

Table S3. NSS as predictor: Comparisons among trajectory groups

|  | Good vs. catch-up | Good vs. Middle | Catch-up vs. Middle | Catch-up vs. Poor | Middle vs. Poor |
| --- | --- | --- | --- | --- | --- |
| SCZ (%) | N.S. | N.S. | N.S. | N.S. | N.S. |
| Affective psychosis (%) |  |  |  |  |  |
| SEX Male (%) | N.S. | N.S. | N.S. | N.S. | N.S. |
| White | N.S. | N.S. | χ2 =4.19;  P= 0.04 | N.S. | N.S. |
| AA |  |  |  |  |  |
| Others |  |  |  |  |  |
| Avg IQ score | t = 3.61;  P=0.003 | t = 2.91;  P=0.005 | N.S. | t = 3.03;  P=0.004 | t = 4.03;  P=0.0002 |
| Patient SES | N.S. | N.S. | N.S. | N.S. | N.S. |
| Parent SES | t = 3.30;  P=0.001 | N.S. | N.S. | N.S. | N.S. |
| Age onset | N.S. | N.S. | N.S. | N.S. | N.S. |
| Substance use Frequency | N.S. | N.S. | N.S. | N.S. | N.S. |
| BPRS-18 items | N.S. | t = -2.85;  P=0.005 | N.S. | N.S. | N.S. |
| HRSM-24 items | N.S. | N.S. | N.S. | N.S. | N.S. |
| WCST-perseverative errors | N.S. | N.S. | N.S. | N.S. | N.S. |
| PMAS-CHILD | N.S. | N.S. | N.S. | N.S. | N.S. |
| PMAS-TOTAL | N.S. | N.S. | N.S. | N.S. | N.S. |
| Duration of Prodromal (wk) | N.S. | N.S. | N.S. | N.S. | N.S. |
| CLUSTER-A | N.S. | N.S. | t = -2.22;  P=0.03 | N.S. | N.S. |
| CLUSTER-B | N.S. | N.S. | N.S. | N.S. | N.S. |
| CLUSTER-C | N.S. | N.S. | t = -2.89;  P=0.008 | N.S. | N.S. |
